# Millisecond-Scale White Matter Dynamics Underlying Visuomotor Integration

**DOI:** 10.1101/2025.03.28.646029

**Authors:** Riyo Ueda, Hiroshi Uda, Keisuke Hatano, Kazuki Sakakura, Naoto Kuroda, Yu Kitazawa, Aya Kanno, Min-Hee Lee, Jeong-Wong Jeong, Aimee F. Luat, Eishi Asano

## Abstract

In the conventional neuropsychological model, nonverbal visuospatial processing is predominantly handled by the right hemisphere, whereas verbal processing occurs in the left, with right-hand responses governed by the left motor cortex. Using intracranial EEG and MRI tractography, we investigated the timing and white matter networks involved in processing nonverbal visuospatial stimuli, forming response decisions, and generating motor outputs. Within 200 ms of stimulus onset, we observed widespread increases in functional connectivity and bidirectional neural flows from visual to association cortices, predominantly in the right hemisphere. Engagement of the right anterior middle frontal gyrus improved response accuracy; however, the accompanying enhancement in intra-hemispheric connectivity delayed response times. In the final 100 ms before right-hand response, functional connectivity and bidirectional communication via the corpus callosum between the right and left motor cortices became prominent. These findings provide millisecond-level support for the established model of hemispheric specialization, while highlighting a trade-off between accuracy and speed governed by the right dorsolateral prefrontal network. They also underscore the critical timing of callosal transmission of response decisions formed in right-hemispheric networks to the left-hemispheric motor system.

**Highlights:**

Neural information propagates through fasciculi during a visuomotor task.
Non-verbal visuospatial analysis is mediated with right-hemispheric dominance.
The right middle frontal gyrus improves response accuracy but delays responses.
Interhemispheric information transfer occurs immediately before motor responses.
This transfer between motor cortices is mediated by the corpus callosum.

## 1. Introduction

Humans constantly engage in visuomotor integration, a process that operates on a millisecond timescale. This process is crucial for activities such as driving a car, navigating crowded urban environments without colliding with pedestrians, and competing in one-on-one sports. Visuomotor integration is also utilized in work settings—for instance, when identifying and removing defective products from a conveyor belt in a factory or inspecting eggs at a poultry farm to check for cracks or dirt and then selecting them by hand. As a result, most people typically use their dominant hand for these tasks (Gilbert and Wysocki, 1992; Peters et al., 2006). Visuomotor integration inherently requires a series of sensory, cognitive, and motor processes, including visual encoding (Hoogenboom et al., 2006; Swettenham et al., 2009; Hermes et al., 2015), feature extraction, decision making, motor planning, and motor execution (Crone et al., 2006; Van Eimeren et al., 2006; Beurze et al., 2007; Ball et al., 2008; Wolynski et al., 2009; Breckel et al., 2011; Kravitz et al., 2011; Brovelli et al., 2017; Combrisson et al., 2017; Nakayama et al., 2022). Although the brain regions involved in visuomotor integration are well documented (as stated below), the millisecond-scale temporal dynamics and the specific white matter pathways that serve as conduits for intra- and inter-hemispheric neural interactions remain understudied.

The current neurobiological model suggests that a bilateral cerebral network originating in the visual cortex is responsible for perceiving and analyzing nonverbal visual stimuli, while the left motor cortex plays a central role in motor execution in right-handed individuals (Hoogenboom et al., 2006; Van Eimeren et al., 2006; Beurze et al., 2007; Swettenham et al., 2009; Wolynski et al., 2009; Breckel et al., 2011; Hermes et al., 2015; Brovelli et al., 2017; Combrisson et al., 2017; Nakayama et al., 2022). Lesion and functional MRI (fMRI) studies point to a dominant role of the right hemisphere in analyzing nonverbal visual stimuli. For instance, lesion studies have shown that visuospatial recognition deficits are more pronounced in patients with right-hemisphere strokes than in those with left-hemisphere strokes (Beis et al., 2004; Habekost, 2007; Urbanski et al., 2011; Wen et al., 2012; Carter et al., 2017). Spatial neglect following right-hemisphere strokes appears to be especially severe when the right arcuate fasciculus and superior longitudinal fasciculus are involved (Urbanski et al, 2011; Carter et al., 2017). In healthy individuals, transcranial magnetic stimulation (TMS) applied to inhibit the right parietal region decreases visuospatial recognition (Mahayana et al., 2014). In addition, fMRI studies in healthy right-handed participants reveal that stronger hemodynamic responses in right-hemispheric association cortices, including the middle-frontal gyrus (MFG), correlate with more accurate performance in tasks requiring nonverbal visuomotor integration (Wen et al., 2012). Single-neuron recordings in non-human primates indicate increased firing rates in the right dorsolateral prefrontal cortex when the animals attend to stimuli during a visuomotor task (Lennert et al., 2011). While it is plausible that visuospatial information generated in the right hemisphere is transferred to the motor network in the left hemisphere for motor execution (Crone et al., 2006; Ball et al., 2008), the timing and the specific white matter pathways involved in this transfer remain uncertain.

Both inter- and intra-hemispheric white matter pathways are potential channels for neural communication with the left-hemispheric motor system, enabling motor responses. In healthy humans, diffusion-weighted imaging (DWI) tractography suggests that the left superior longitudinal fasciculus and the left inferior fronto-occipital fasciculus are among the key intra-hemispheric pathways directly connecting the left visual and motor networks (Yap et al, 2013; Karolis et al, 2017; Maier-Hein et al., 2017). Intracranial studies in patients with focal epilepsy show that single-pulse electrical stimulation of the visual areas elicits cortico-cortical evoked potentials (CCEPs) within 40 ms in the ipsilateral frontal lobe, including the precentral gyrus— evidence that intra-hemispheric fasciculi can directly convey neural information between visual and motor networks (Keller et al., 2014; Oane et al., 2020; Sugiura et al., 2020). Further, studies in non-human primates indicate that, during tasks requiring visuomotor integration, neural information is transferred from the left posterior head region to temporal and frontal lobe regions, including the motor system (Lee et al., 2003; O’Leary et al., 2006; Ledberg et al., 2007). In contrast, tractography studies show that callosal fibers directly connect homotopic motor networks in the right and left hemispheres (Hofer and Frahm, 2006; Shen et al., 2015), while single-pulse electrical stimulation of the motor area in one hemisphere can elicit CCEPs in the corresponding region of the contralateral hemisphere within 40 ms (Terada et al., 2008; Matsumoto et al., 2017; Mitsuhashi et al., 2020).

In the present study, we have constructed a dynamic tractography map (Silverstein et al., 2020; Sonoda et al., 2021) that visualizes intra- and inter-hemispheric neural interactions through specific white matter pathways during a visuomotor task. We quantified task-related neural engagement in regions of interest (ROIs) by measuring high-gamma activity (70–110 Hz) in intracranial EEG (iEEG) recordings. High-gamma activity has been used to assess cortical activation and deactivation with a temporal resolution on the order of tens of milliseconds (Crone et al., 2006; Hoogenboom et al., 2006; Ball et al., 2008; Swettenham et al., 2009; Hermes et al., 2015). Its signal fidelity, when recorded intracranially, is over 100 times greater than that of scalp EEG (Ball et al., 2009). Moreover, high-gamma amplitudes correlate closely with single-neuron firing rates (Ray et al., 2008; Ray et al., 2011; Leszczyński et al., 2020), hemodynamic responses in fMRI (Mukamel et al., 2005; Nir et al., 2007), and glucose metabolism in positron emission tomography (Nishida et al., 2008). We evaluated neural interactions through specific white matter pathways using functional connectivity and directional neural information flow analyses. In this study, we defined functional connectivity enhancement as a significant, simultaneous, and sustained increase in high-gamma amplitude in ROIs directly connected by a DWI-based white matter tract (Kitazawa et al, 2023; Ono et al, 2023; Sakakura et al., 2024; Ueda et al., 2024). We have previously shown that cortical sites exhibiting simultaneous task-related high-gamma augmentation are likely to have underlying structural and effective connectivity, as evidenced by DWI tractography and CCEPs elicited by single-pulse electrical stimulation (Sonoda et al., 2021). We also estimated neural information flow using transfer entropy analysis (Ito et al., 2011; Firestone et al., 2023). We considered a facilitatory information flow to occur from one ROI to another if (1) an increase in high-gamma amplitude at the first ROI predicted a subsequent increase at the second ROI, and (2) the ROIs were directly connected via a white matter tract. Building on fMRI findings that increased hemodynamic responses in right-hemispheric association cortices correlate with better visuomotor performance (Wen et al., 2012), we hypothesized that an increase in right-hemispheric high-gamma amplitudes would be most strongly associated with enhanced behavioral performance, measured by response accuracy and response time. If confirmed, this would clarify the predominantly right-hemispheric processes underlying visuomotor integration. We also tested the hypothesis that neural communication from the right to the left motor system occurs via callosal fibers after the period in which right-hemispheric high-gamma amplitudes influence behavioral performance but before the motor response. If confirmed, this would suggest that the corpus callosum conveys mental representations of response decisions formed in right-hemispheric networks to the left-hemispheric motor system.

## 2. Methods

### 2.1. Participants

The present study included eight participants (**Table 1**) who fulfilled the following eligibility criteria. The inclusion criteria were: (i) patients with focal epilepsy who underwent extraoperative iEEG recording as part of the clinical management of drug-resistant seizures at Children’s Hospital of Michigan between September 2017 and December 2019; (ii) patients able to complete five sessions of *Speed Match* - a visuomotor task on the Lumosity platform (https://www.lumosity.com/; Lumos Labs, Inc, San Francisco, CA), during the iEEG recording. Participants were excluded if they had (i) massive brain malformations that deformed the central, lateral, or calcarine sulcus (Kitazawa et al., 2023), (ii) a history of previous resective epilepsy surgery, (iii) hemiparesis, (iv) visual field deficit on confrontation, or (v) hearing deficit.

**Table 1.** Patient demographics. CLB: Clobazam. LCM: Lacosamide. LEV: Levetiracetam. LTG: Lamotrigine. OXC: Oxcarbazepine. TPM: Topiramate. VPA: Valproate. ZNS: Zonisamide. F: Frontal. O: Occipital. P: Parietal. T: Temporal. NA: Not available because seizure events did not occur during the intracranial EEG recording.

| Patient number | Age (years) | Sex | Sampled hemisphere | Number of analyzed electrodes | Antiepileptic drugs | Age of epilepsy onset (years) | Seizure onset zone | MRI finding |
| --- | --- | --- | --- | --- | --- | --- | --- | --- |
| 1 | 11 | Male | Left | 74 | OXC | 6 | T | Tumor |
| 2 | 16 | Male | Left | 89 | LEV | 9 | T | Tumor |
| 3 | 20 | Male | Right | 105 | VPA, OXC | 6 | F | Nonlesional |
| 4 | 9 | Male | Left | 64 | OXC | 9 | T | Tumor |
| 5 | 16 | Female | Right | 114 | CLB, TPM | 12 | NA | Nonlesional |
| 6 | 14 | Female | Right | 79 | LCM, LTG | 5 | Insula | Tumor |
| 7 | 15 | Female | Right | 106 | LTG, ZNS | 6 | P | Nonlesional |
| 8 | 19 | Female | Left & Right | 95 | OXC | 4 | F | Nonlesional |

All participants were right-handed, and none exhibited a congenital, left-hemispheric neocortical MRI lesion associated with left-handedness. Therefore, all patients were suggested to have essential language areas located in the left hemisphere (Rasmussen and Milner, 1977; Akanuma et al., 2003; Möddel et al., 2009). We had a comprehensive discussion on the justification and reliability of estimating the language-dominant hemisphere through anatomical imaging and handedness (Sonoda et al., 2022). Indeed, electrical stimulation mapping identified left-hemispheric language sites in all four patients who underwent intracranial electrode sampling mainly from the left hemisphere (i.e., patients 1, 2, 4, and 8). No such sites were detected in the right hemisphere of any patient.

### 2.2. Intracranial EEG (iEEG) and MRI data acquisition

The data acquisition framework for iEEG and MRI data was the same as previously reported (Nakai et al., 2017; Johnson et al., 2018; 2022; Mitsuhashi et al., 2022). Platinum disk electrodes with a 10 mm center-to-center distance were surgically placed on the pial surface of the brain (**Figure 1**). The number and configuration of intracranial electrodes were based purely on clinical needs to localize the boundary between the epileptogenic zone and eloquent areas, and no extra electrodes were implanted for research purposes. Bedside iEEG recording was performed using a Nihon Kohden Neurofax Digital System (Nihon Kohden America Inc, Foothill Ranch, CA, USA) with a sampling rate of 1,000 Hz and an amplifier bandpass filter of 0.016-300 Hz. The exact timing of stimulus onset, as well as participants’ tap responses during the visuomotor task, were integrated into the iEEG acquisition system via the DC input (Mitsuhashi et al., 2022). A total of 637 electrode sites were utilized for subsequent iEEG analysis, after excluding those located in the seizure onset zone (Asano et al., 2009), spiking zone (Kural et al., 2020), or structural lesions, along with those affected by artifacts.

**Figure 1.**
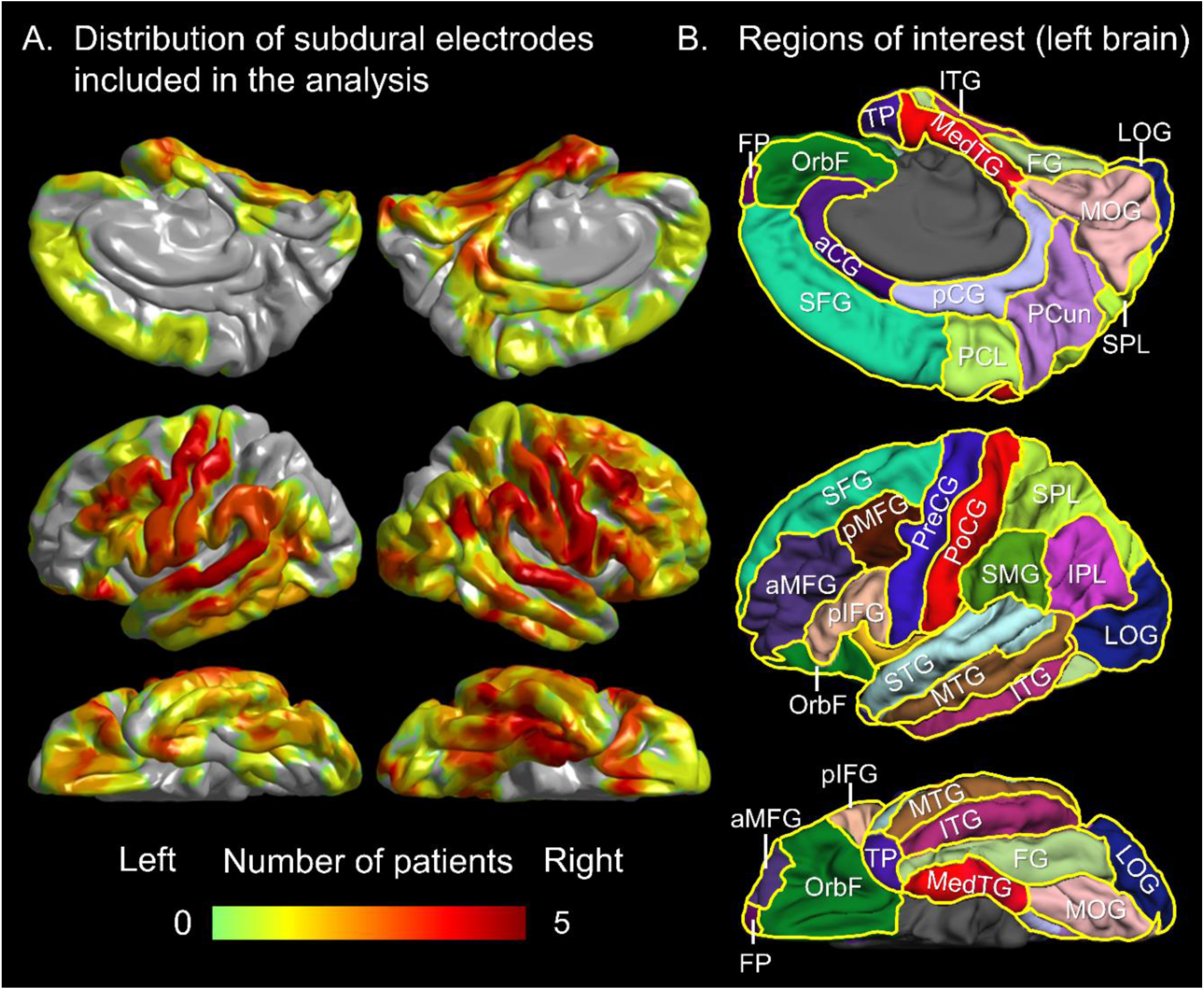
Distribution of subdural electrode sites across regions of interest (ROIs). (A) The pooled distribution of electrode sites from eight patients. (B) ROI locations in left hemisphere. aMFG and pMFG: anterior and posterior middle-frontal gyri. FG: fusiform gyrus. IPL: inferior-parietal lobule. ITG: inferior-temporal gyrus. LOG: lateral-occipital gyrus. MedTG: medial-temporal gyrus (summation of entorhinal and parahippocampal gyri). MOG: medial-occipital gyrus (summation of cuneus and lingual gyri). MTG: middle-temporal gyrus. pIFG: posterior inferior-frontal gyrus (summation of pars opercularis and pars triangularis). PoCG: postcentral gyrus. PreCG: precentral gyrus. SFG: superior-frontal gyrus. SMG: supramarginal gyrus. STG: superior-temporal gyrus.

Before intracranial electrode placement, we acquired 3-tesla MRI scans, which included T1-weighted spoiled gradient-recalled echo and fluid-attenuated inversion recovery sequences (Nakai et al., 2017). We used FieldTrip (http://www.fieldtriptoolbox.org) to create a 3D MRI surface image, where electrode locations were defined directly on the brain surface using post-implant CT images (Stolk et al., 2018). We next used FreeSurfer (http://surfer.nmr.mgh.harvard.edu) to normalize each electrode site to a Montreal Neurological Institute (MNI)-standard brain coordinate for group-level visualization and analysis. We then divided the cerebral cortex of each hemisphere into 21 ROIs based on the Desikan parcellation (Desikan et al., 2006) and included 30 ROIs symmetrically located in both hemispheres (15 ROIs in each hemisphere), each containing at least three electrodes, in our group-level statistical analyses (**Figure 1**).

All aforementioned ROIs (15 per hemisphere) were included in the ROI-based analysis, with each containing at least three electrodes. **Table S1** presents the number of electrode sites within each ROI. The ROI-based analysis was not conducted in the following regions due to an insufficient sample (**Table S2**). aCG: anterior-cingulate gyrus (summation of rostral and caudal anterior-cingulate gyri). FP: frontal pole. OrbF: orbitofrontal gyrus (summation of pars orbitalis and medial-and lateral-orbitofrontal gyri). pCG: posterior-cingulate gyrus (summation of posterior and isthmus cingulate gyri). PCL: paracentral lobule. PCun: precuneus gyrus. SPL: superior-parietal lobule.

### 2.3. Visuomotor task

Patients completed the *Speed Match* in a quiet room, during their interictal state and at least two hours apart from habitual seizure events (Figure 2A; **Video S1**). None of the patients had previously performed this visuomotor task, and all patients were given a brief tutorial to understand the rules prior the first session. Patients were comfortably positioned on bed, and used an iPad (screen display width: 14.7 cm; length: 19.6 cm; Apple Inc., Cupertino, CA) to play five consecutive sessions, each lasting 45 seconds. During a given trial, one of the five symbols was presented. Each patient was instructed to tap “Yes” with their right index finger if an incoming symbol matched the previous one, and “No” if it did not. The response time for a given trial was defined as the period between stimulus onset and response onset.

**Figure 2.**
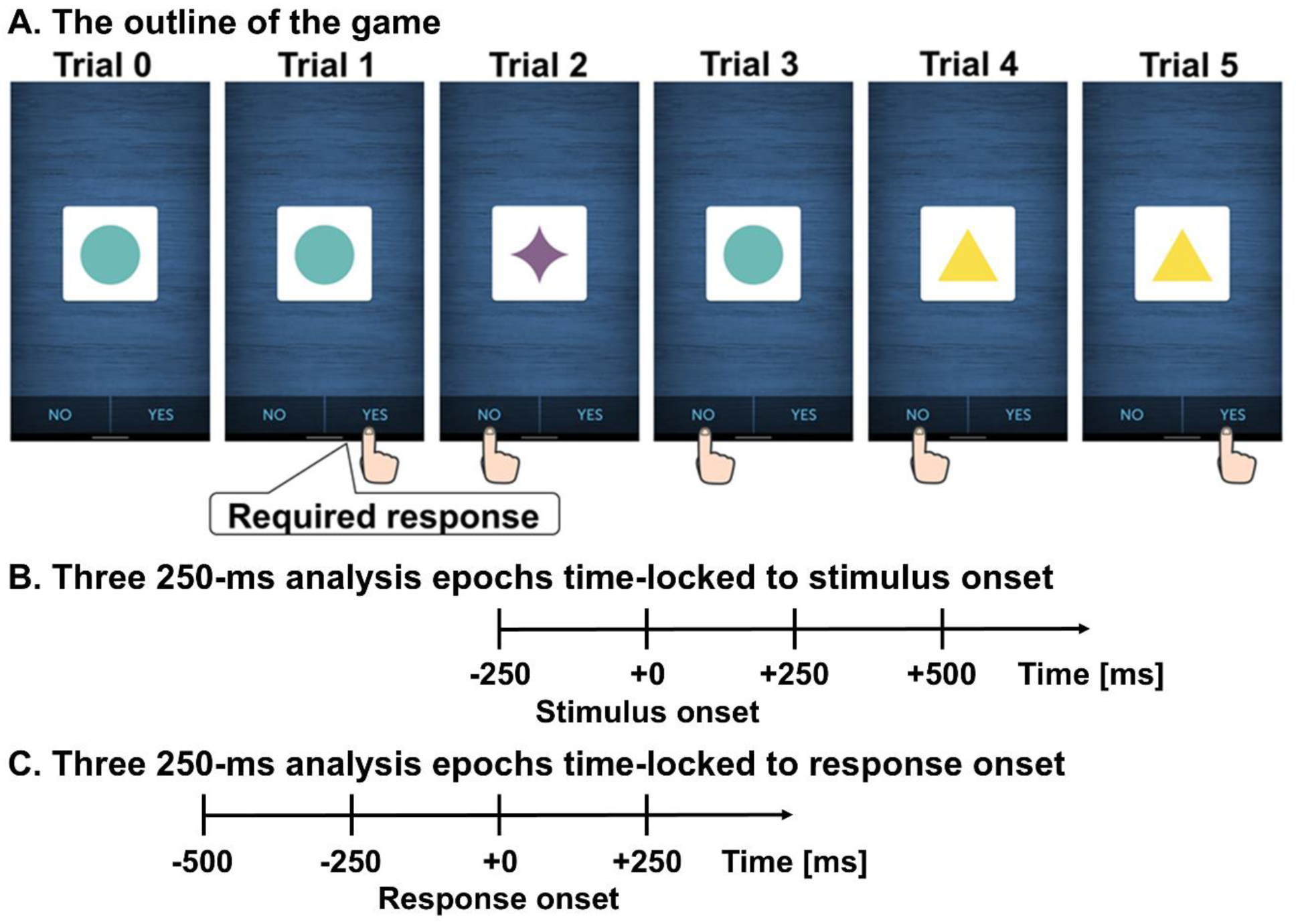
*Speed Match*: a visuomotor task. (A) Overview of the game: During a given trial, one of the five symbols was presented (**Video S1**). Each patient was instructed to tap “Yes” with their right index finger if an incoming symbol matched the previous one, and “No” if it did not. The response time for a given trial was defined as the period between stimulus onset and response onset. (B) Three 250-ms analysis epochs time-locked to stimulus onset, defined as the display of a symbol. (C) Three 250-ms analysis epochs time-locked to response onset, marked by the finger tapping.

### 2.4. Time-frequency analysis

At each electrode channel, we performed time-frequency analysis on iEEG signals referenced to the common average, as previously reported (Nakai et al., 2019; Mitsuhashi et al., 2022; Kitazawa et al., 2023; Ono et al., 2023). The complex demodulation method incorporated in the BESA EEG Analysis Package (BESA GmbH, Gräfelfing, Germany; Papp and Ktonas, 1977; Hoechstetter et al., 2004) transformed iEEG signals into 5-ms/10-Hz time-frequency bins for assessment of high-gamma amplitude70-110 Hz during the 750-ms analysis windows (Figure 2B**, 2C**). The complex modulation was done by multiplying the time-domain iEEG signal with a complex exponential, followed by a band-pass filter. Because it employed a Gaussian-shaped low-pass finite impulse response filter, this complex demodulation method is equivalent to a Gabor transformation. The time-frequency resolution for high-gamma measurement was ±7.9 ms and ±14.2 Hz (defined as the 50% power drop of the finite impulse response filter). To visualize the spatiotemporal dynamics of task-related high-gamma modulations, we calculated the percent change in high-gamma amplitude (where amplitude is a measure proportional to the square root of power) for each electrode site, in comparison to the mean of the entire 750-ms analysis period, containing 150 time bins. We then generated a video visualizing the percentage change in high-gamma amplitude on a FreeSurfer standard pial surface image, with interpolation applied within 10 mm of each electrode center (Sakakura et al. 2022; **Video S2**).

### 2.5. Assessment of task-related high-gamma amplitude modulations

At each ROI, high-gamma augmentation and attenuation were assessed using a mixed model analysis applied to each 5-ms time bin across the 750-ms analysis epoch (150 time bins). The dependent variable was high-gamma amplitude modulation (% change) at each time bin for each trial. The fixed effect was the intercept, while the random effects included electrode and patient, allowing the intercept to vary across electrodes and patients. We deemed high-gamma augmentation significant when the lower limit of the 95% confidence interval (95% CI) of the intercept exceeded zero for at least 20 ms (4 consecutive time bins). Similarly, high-gamma attenuation was considered significant when the upper limit of the 95% CI of the intercept fell below zero for at least 20 ms. These thresholds correspond to a chance probability (Type I error) of approximately 0.0919%, calculated as 0.05^4^ × (150 - 3) × 100.

### 2.6. Structural connectivity

To delineate structural white matter streamlines, we utilized open-source DWI data from 1,065 healthy participants (Yeh et al., 2018; http://brain.labsolver.org/diffusion-mri-templates/hcp-842-hcp-1021); a method previously reported in our work (Mitsuhashi et al., 2021; Sonoda et al., 2021; Mitsuhashi et al., 2022; Kitazawa et al., 2023; Ono et al., 2023; Sakakura et al., 2023). The DSI Studio script (http://dsi-studio.labsolver.org/) identified tractography streamlines directly connecting a pair of cortical ROIs within the MNI standard space. The fiber tracking parameters used were a quantitative anisotropy threshold of 0.05, a maximum turning angle of 70° and a streamline length of 20 to 250 mm. A board-certified neurosurgeon (K.S. and H.U.) excluded streamlines involving the brainstem, basal ganglia, thalamus, or cerebrospinal fluid space from further analysis.

### 2.7. Functional connectivity

Because iEEG can only be sampled from limited areas in a given patient, a group-level summary measure derived from a different subset of patients (such as mean high-gamma amplitudes at each ROI) is commonly used in iEEG studies to estimate the neural communication dynamics shared by the general population (Kunii et al., 2013; Burke et al., 2014; Avanzini et al., 2016; Solomon et al., 2017; Kanno et al., 2025). An appropriate analogy for this group-level analysis is conducting a cross-sectional study to infer temporal effects on neural measures when a longitudinal design with the same participants is impractical. We validated the use of group mean high-gamma amplitudes for computing functional connectivity enhancements by demonstrating a significant association (Spearman’s correlation coefficient range: +0.54 to +0.82; p-value range: 2.5 × 10⁻⁵ to 6.6 × 10⁻¹⁴) with specific sensory, motor, and cognitive manifestations elicited by electrical stimulation mapping (Kanno et al., 2025). We acknowledge that increasing the number of electrode sites used to compute the group mean reduces the influence of any single electrode site. For transparency, we have provided the number of electrode sites included in our analysis in **Table S1**.

Using a method similar to those reported previously (Kitazawa et al., 2023; Ono et al., 2023), we generated a dynamic tractography video that visualizes the spatiotemporal characteristics of functional connectivity through white matter tracts. We declared functional connectivity to be enhanced (or diminished) between a pair of ROIs only if (i) both cortical ROIs exhibited significant, simultaneous, and sustained (at least 20 ms) augmentation (or attenuation) in high-gamma amplitude, and (ii) they were connected by direct DWI streamlines. When significant high-gamma augmentation occurred in χ% of time of the 250 time bins during 1,250 ms analysis period (from 500 ms pre-response onset to 500 ms post-stimulus onset), the chance probability of observing co-occurring high-gamma augmentation in at least one pair among 30 ROIs was 1 – (1 – ((χ/100)^2^)^4^)^435^. Given that 30 ROIs were analyzed, there were 435 unique ROI pairs. By color-coding the relevant white matter streamlines, we depicted enhancements and diminutions in functional connectivity at 5-ms intervals throughout the 750-ms analysis period.

### 2.8. Neural information flows

We quantified neural information flows using a transfer entropy-based effective connectivity analysis (Firestone et al., 2023). Computational analysis of neural information flows inherently requires a certain length of data points for reliable analysis (Korzeniewska et al., 2011; Johnson et al., 2018). We conducted analyses on each of three sequential 250-ms, spanning from 250 ms before stimulus onset to 500 ms after stimulus onset (Figure 2B), as well as another set of three sequential 250-ms epochs, spanning from 500 ms before response onset to 250 ms after response onset (Figure 2C). We declared that facilitatory information flow took place from ROI1 to ROI2 during a given 250-ms analysis epoch if all of the following three criteria were met: [1] ROI1 and ROI2 were connected by a direct white matter tract on DWI tractography. [2] Functional connectivity enhancement was noted between ROI1 and ROI2 during a given 250-ms analysis epoch of interest. [3] The transfer entropy program (Ito et al., 2011) implemented in Matlab R2020a (MathWorks, Natick, MA, USA) indicated that, within the 250-ms epoch, a binary time series of high-gamma amplitude data at ROI1 predicted the data at ROI2. Each 5-ms time bin was assigned a value of 1 if significant high-gamma augmentation was noted in a given ROI; otherwise, it was assigned a value of 0. Because transfer entropy inherently measures conditional mutual information (Schreiber, 2000) in “bits,” binarizing the continuous high-gamma amplitude data was necessary to make it compatible with the equation. The mean strength of information flows with all intra- and inter-hemispheric ROIs, both efferent (originating from the ROI) and afferent (converging on the ROI), was then computed for each ROI.

We likewise computed the strength of suppressive neural information flows between pairs of ROIs exhibiting functional connectivity diminution. For each 5-ms time bin, a value of 1 was assigned if significant high-gamma attenuation was observed in the given ROI; otherwise, a value of 0 was assigned.

### 2.9. Assessment of neural information flows at the hemispheric level

We determined whether the intensity of facilitatory information flows within (i.e., left to left; right to right) and between hemispheres (i.e., left to right; right to left) differed across three 250-ms analysis epochs around stimulus onset (Figure 2B). To this end, we employed Friedman’s two-way ANOVA by rank test. Subsequently, we conducted the Wilcoxon signed-rank test as a post-hoc analysis to compare the flows between two 250-ms analysis epochs. A Bonferroni correction was applied to account for three pairwise comparisons, setting the threshold for statistical significance at an uncorrected two-sided p-value of less than 0.0167. We specifically hypothesized that facilitatory information flows within and toward the right hemisphere would be higher after stimulus onset compared to before. The same analysis was repeated to assess whether suppressive information flows varied across the three 250-ms analysis epochs. Additionally, we conducted the analysis using epochs around response onset (Figure 2C).

### 2.10. Assessment of inter-hemispheric neural information flows at the region-to-region level

We subsequently determined the temporal dynamics of inter-hemispheric facilitatory information flows, rated by transfer entropy, between the homotopic ROIs showing functional connectivity enhancements. To this end, we calculated transfer entropy values for each 250 ms epoch, sliding in 10 ms increments during the analysis epoch time-locked to stimulus as well as response onset. We employed Spearman’s rank correlation to examine whether inter-hemispheric facilitatory information flows increased or decreased as the stimulus presentation approached and progressed thereafter, and likewise as the response approached and progressed thereafter. We specifically predicted that inter-hemispheric facilitatory information flows toward the left precentral gyrus would increase as the response approached.

### 2.11. Relationship between high-gamma modulation and behavioral performance

Logistic mixed model analysis was conducted to determine whether enhanced high-gamma amplitudes at a given ROI were associated with improved response accuracy, independent of stimulus properties. The dependent variable was response accuracy, with a value of 1 indicating a correct response. The fixed effect variables included: [a] ‘different symbol’: whether the presented symbol differed from that in the prior trial, [b] ‘task switch’: whether the required response differed from that in the prior trial, [c] ‘task familiarity’: a value ranging from 0 to the total number of trials in each session for each patient, representing the degree of task familiarity, [d] ‘prior failure’: whether a wrong response was made in the prior trial, and [e] ‘mean high-gamma’: the mean during a given 50-ms analysis epoch at a specific electrode. Patient and electrode were treated as random factors, allowing the intercept to vary across electrodes and patients. A Bonferroni correction was applied to account for 10 repeated models across 50-ms analysis epochs between stimulus onset and 500 ms after stimulus onset, setting the threshold for statistical significance at an uncorrected two-sided p-value of less than 0.005.

Linear mixed model analysis was subsequently performed to investigate whether enhanced high-gamma amplitudes at a given ROI were associated with altered response time. In this case, the dependent variable was response time for each trial, while the fixed and random effect variables remained the same as above. Similarly, an uncorrected two-sided p-value of less than 0.005 was used as the threshold for significance. All statistical analyses were performed using IBM SPSS Statistics 25 (IBM Corp., Chicago, IL, USA).

### 2.12. Ethics statement

The Institutional Review Board at Wayne State University approved this study. We obtained written informed consent from patients 18 years or older or the legal guardians of patients younger than 18. We also obtained written assent from pediatric patients aged 13 years or older.

### 2.13. Data availability

The iEEG data is available at https://openneuro.org/datasets/ds005931 (doi:10.18112/openneuro.ds005931.v1.0.0).

### 2.14. Code availability

The analysis codes are available at http://github.com/rio-k12b1r1/Speed_Match (doi: 10.5281/zenodo.8475).

## 3. Results

### 3.1. Behavioral observations

A given patient completed an average of 184 trials (range: 117–232). The average response time was 1,023 ms (range: 744–1,625 ms). A mixed model analysis, incorporating [a] different symbol, [b] task switch, [c] task familiarity, [d] prior failure, and [e] patient age as fixed effects, with patient as a random effect, identified a stimulus property significantly associated with response time. Specifically, a different symbol was independently associated with prolonged response time (mixed model coefficient: +276 ms; 95% CI: +216 to +336; t-value: +9.0; p-value: 7.3 × 10^-19^). Task familiarity was independently associated with shortened response time (mixed model coefficient: −2 ms/trial; 95% CI: −3 to −1; t-value: −7.4; p-value: 2.0 × 10^-13^). Likewise, prior failure was independently associated with prolonged response time (mixed model coefficient: +306 ms; 95% CI: +205 to +407; t-value: +5.9; p-value: 3.6 × 10^-9^). No other fixed effects were significantly associated with response time (p-values: 0.16 to 0.28; **Table S3**).

The correct response rate averaged 87.9% (range: 74.7%–94.5%). A logistic mixed model analysis, incorporating the aforementioned fixed and random effect factors, showed that prior failure was associated with a lower likelihood of a successful trial (odds ratio: 0.402; 95% CI: 0.248 to 0.651; t-value: −3.7; p-value: 2.1×10^-4^), and different symbol was associated with a lower likelihood of a successful trial (odds ratio: 0.647; 95% CI: 0.442 to 0.948; t-value: −2.2; p-value: 0.026) as well. No other fixed effects were significantly associated with response accuracy (p-values: 0.09 to 0.93; **Table S4**).

### 3.2. Dynamics of cortical high-gamma modulations

**Video S2** presents an overview of task-related high-gamma modulations. Following stimulus onset, the occipital and fusiform regions exhibited high-gamma augmentation, whereas the left precentral gyrus showed high-gamma attenuation. By 250 ms before responses, the right-hemispheric association cortices, including the anterior middle-frontal gyrus (aMFG), exhibited high-gamma augmentation. As the response approached, high-gamma augmentation in the right and left precentral gyri intensified. Around responses, the occipital and fusiform regions exhibited high-gamma attenuation.

Mixed model analysis identified the precise timing of high-gamma modulations. Specifically, high-gamma augmentation in the left lateral occipital gyrus was maximized at 250 ms post-stimulus onset, based on the t-value (mixed model coefficient: +9.8%; 95% CI: +5.8 to +13.8; t-value: +5.5; p-value: 3.8 × 10⁻⁴; DF: 9.0). High-gamma augmentation in the right lateral occipital gyrus was maximized at 235 ms (+10.8%; 95% CI: +4.4 to +17.2; t-value: +3.5; p-value: 0.003; DF: 17.1). High-gamma augmentation in the right fusiform gyrus was maximized at 290 ms (+6.7%; 95% CI: +3.5 to +9.9; t-value: +4.5; p-value: 3.2 × 10⁻⁴; DF: 17.1).

High-gamma augmentation in the right aMFG became significant at 240 ms prior to response, and it was maximized at 25 ms pre-response onset (+4.8%; 95% CI: +2.7 to +6.9; t-value: +4.7; p-value: 6.9 × 10^-5^; DF: 27.5). High gamma augmentation in the right precentral gyrus was maximized at 25 ms pre-response onset (+4.2%; 95% CI: +1.5 to +6.8; t-value: +3.1; p-value: 0.003; DF: 46.1). High gamma augmentation in the left precentral gyrus became significant at 125 ms prior to response and was maximized at 75 ms after response (+6.1%; 95% CI: +4.1 to +8.0; t-value: +6.3; p-value: 2.1 × 10^-7^; DF: 39.9).

### 3.3. Statistical validity of the reported functional connectivity modulations

Significant high-gamma augmentation and attenuation were noted in 11.97% and 10.93% of the time bins, respectively, on average across ROIs. The chance probability of simultaneous significant high-gamma augmentation and attenuation at two or more ROIs lasting for ≥40 ms was 0.0018% and 0.00089%, respectively. In other words, the functional connectivity enhancement and diminution, reported below, had very low Type I error rates.

### 3.3. Overview of functional connectivity and neural information flow

Readers are encouraged to view **Videos S3** and **S4** before proceeding. **Video S3** provides a comprehensive view of functional connectivity modulations and neural information flow between anatomical ROI pairs across time windows. **Video S4** illustrates the specific white matter pathways supporting functional connectivity, along with the direction and strength of information flow.

During the first 250 ms following stimulus onset, functional connectivity increased between the occipital and temporal regions via the inferior longitudinal fasciculi (Figure 3B). Simultaneously, neural information propagated from the lateral occipital gyrus to the fusiform and medial temporal regions, as well as from the left to right occipital lobes. Over the next 250 ms, functional connectivity via the inferior longitudinal fasciculus and local u-fibers became predominantly right-hemispheric between the occipital and temporal regions (Figure 3B). During the 500 ms following stimulus onset, functional connectivity diminution was noted between the precentral and postcentral gyri.

**Figure 3.**
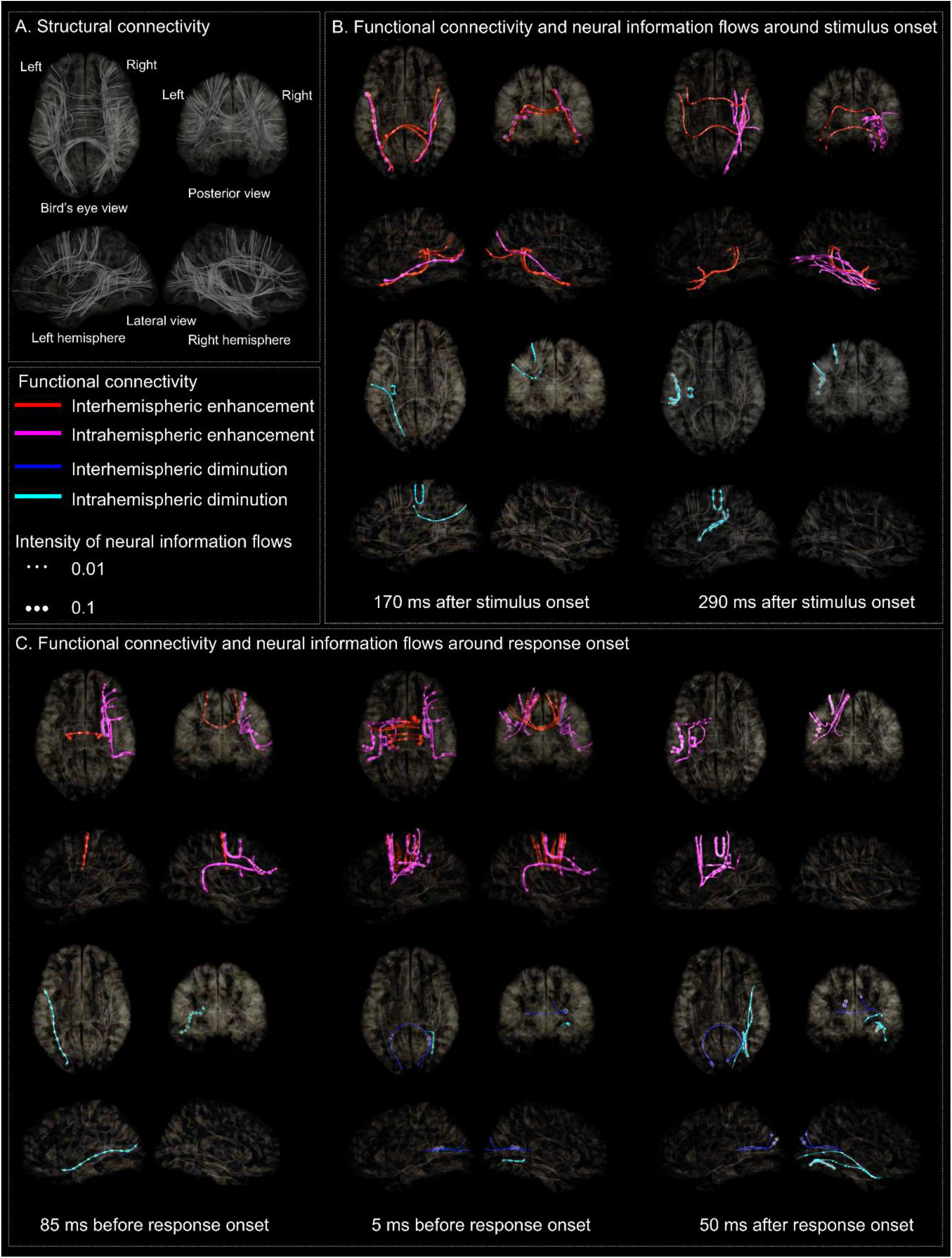
Functional connectivity and neural information flows. (A) Structural connectivity. (B) and (C) show functional connectivity and neural information flows around stimulus onset and response onset, respectively. Color-coded streamlines represent different types of functional connectivity modulations. **Video S4** visualizes the direction of neural information flow using moving spheres.

During the 500 ms prior to response, functional connectivity was enhanced across four lobes, mainly within the right hemisphere via the superior longitudinal and arcuate fasciculi (Figure 3C). As the response onset approached, functional connectivity increased between the right MFG and precentral gyrus via local u-fibers. In the 100 ms before response, inter-hemispheric functional connectivity increased between the right and left precentral gyri via callosal fibers, with bidirectional information transfer (Figure 3C). Following response onset, functional connectivity was enhanced between the left precentral and postcentral gyri via local u-fibers, accompanied by bidirectional information flows, whereas functional connectivity within the right hemisphere decreased.

### 3.4. Statistical assessment of neural information flows at the hemispheric level

The results indicate that intra-hemispheric neural communication within the right hemisphere is most active during the first 250 ms after stimulus onset and least active immediately after the response. Specifically, right-hemispheric facilitatory neural information flow was higher during the first 250 ms after stimulus onset compared to the 250–500 ms post-stimulus period (p-value: 0.002 on Wilcoxon signed-rank test; effect size r: 0.35; Figure 4A). Facilitatory flow during the 250–500 ms post-stimulus period was higher than during the 250 ms preceding stimulus onset (p-value: 6.5 × 10^-4^; r: 0.54). Conversely, right-hemispheric suppressive neural information flow was less intense during the first 250 ms after stimulus onset compared to the 250–500 ms post-stimulus period (p-value: 6.5 × 10^-4^; r: 0.55; Figure 4B) and the 250 ms preceding stimulus onset (p-value: 4.4 × 10^-4^; r: 0.53).

**Figure 4.**
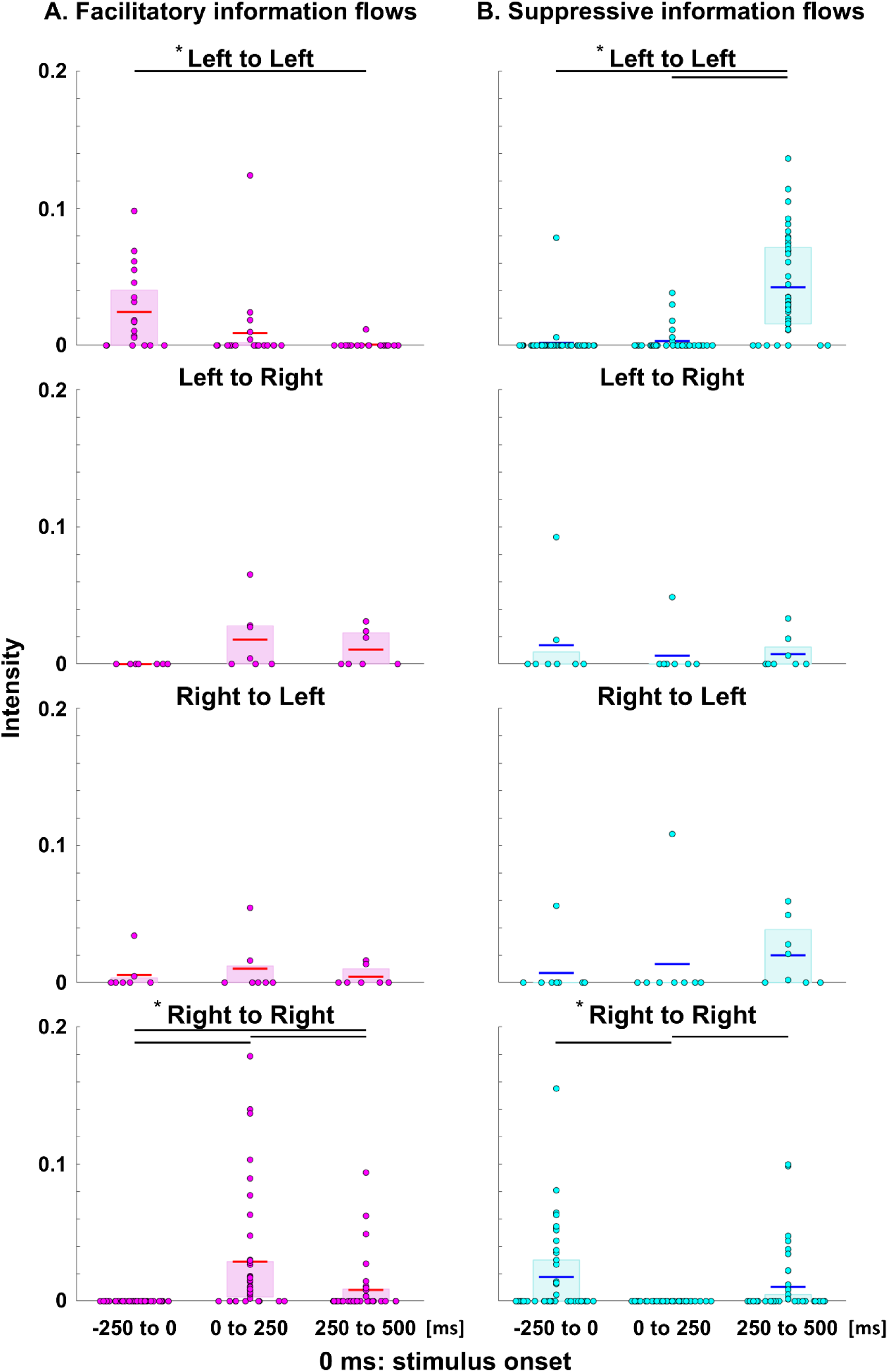
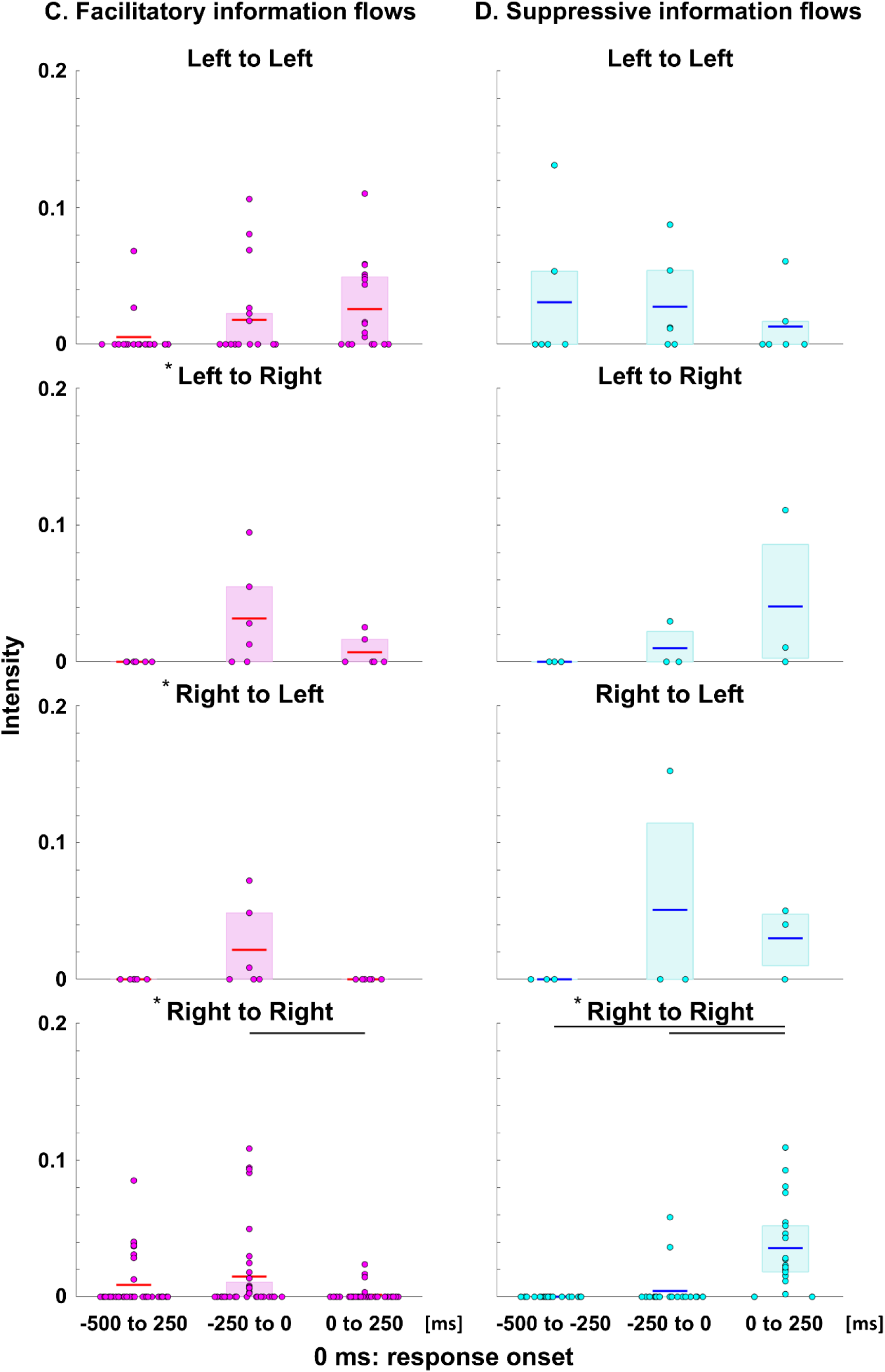
Statistical assessment of neural information flow at the hemispheric level. (A) Box plots depict the intensity of facilitatory neural information flows measured at 15 ROIs during three 250-ms time windows around stimulus onset. (B) Suppressive flows around stimulus onset. (C) Facilitatory flows around response. (D) Suppressive flows around response. The red bars in (A) and (C), and as well as the blue bars in (B) and (D) indicate the mean intensity of neural information flows. An asterisk denotes a significant difference in the intensity across the three-time windows, assessed using Friedman’s two-way ANOVA by ranks. A horizontal black bar indicates a significant pairwise difference, determined by the Wilcoxon signed-rank test.

Right-hemispheric facilitatory neural information flow during the first 250 ms after response onset was lower than during the 250 ms preceding response onset (p-value: 2.0 × 10^-4^; r: 0.59; Figure 4C). Conversely, right-hemispheric suppressive neural information flow during the first 250 ms after response onset was stronger than during the 250–500 ms pre-response period (p-value: 8.9 × 10^-5^; r: 0.84; Figure 4D) and the 250 ms immediately prior to response (p-value: 0.002; r: 0.66).

The results also indicate that inter-hemispheric neural communication within the left hemisphere is most active around response. Left-hemispheric facilitatory neural information flow was higher during the 250 ms preceding stimulus onset (i.e., immediately after the response) compared to the 250–500 ms post-stimulus period (p-value: 0.001; r: 0.71; Figure 4A). Conversely, left-hemispheric suppressive flow was lower during the 250 ms preceding stimulus onset compared to the 250–500 ms post-stimulus period (p-value: 5.9 × 10^-7^; r: 0.75; Figure 4B).

### 3.5. Statistical assessment of inter-hemispheric neural information flows at the region-to-region level

Figure 5 shows the temporal evolution of inter-hemispheric, facilitatory neural information flows among ROIs with enhanced functional connectivity. Toward 250 ms post-stimulus onset, neural information flows increased from the left lateral occipital gyrus to the right (Spearman’s correlation coefficient: +0.92; p-value: 1.7 × 10^-11^) and also from the right lateral occipital gyrus to the left (Spearman’s correlation coefficient: +0.60; p-value: 0.001). Following this, neural information flows from the right to left lateral occipital gyrus decreased (Spearman’s correlation coefficient: −0.54; p-value: 0.005).

**Figure 5.**
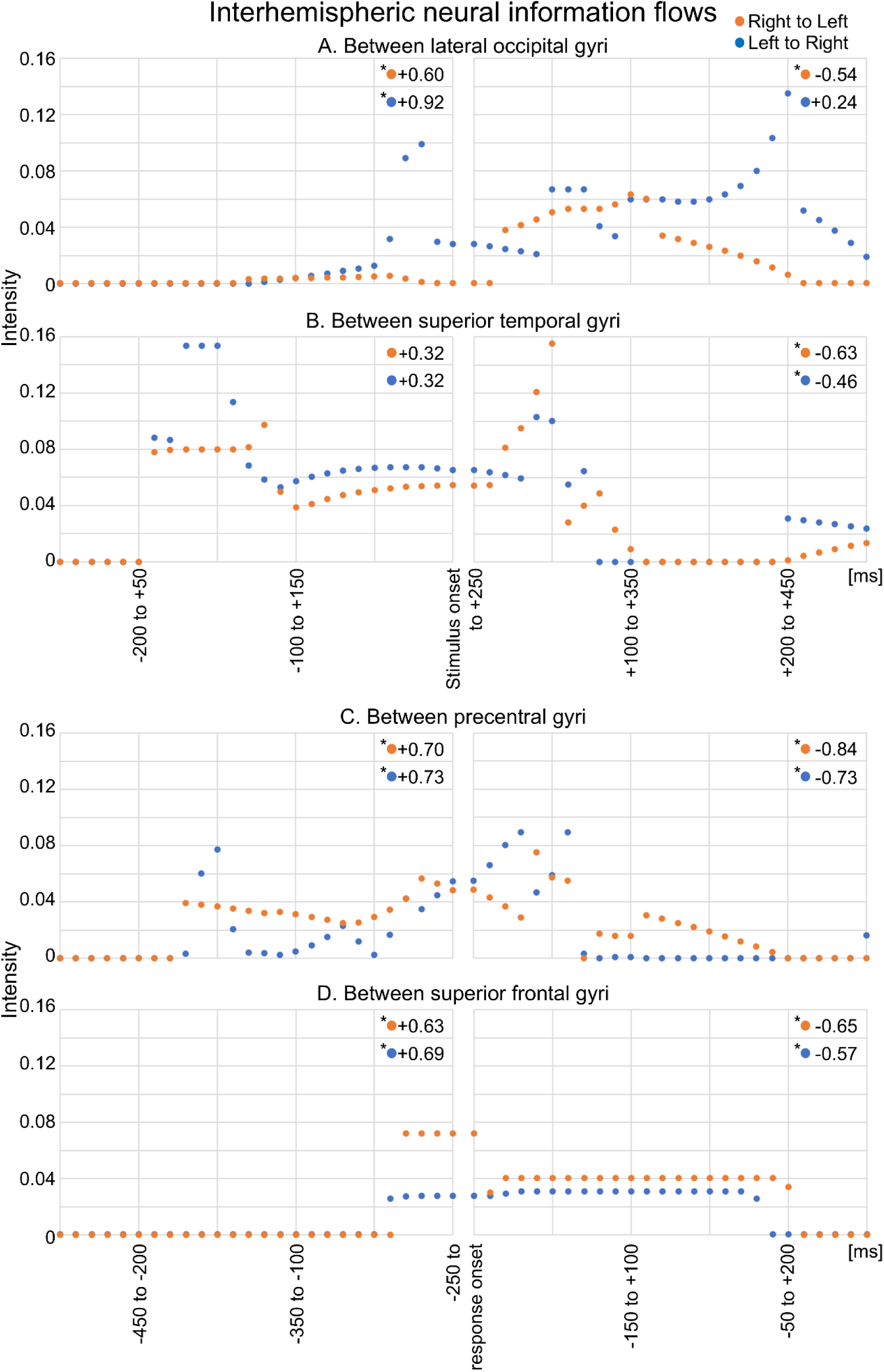
Statistical assessment of inter-hemispheric neural information flow at the region-to-region level. (A and B) The plots show changes in the intensity of inter-hemispheric facilitatory neural information flow around stimulus and response, respectively. The orange plot represents flow from the right to the left hemisphere, while the blue plot represents flow from the left to the right hemisphere. An asterisk denotes a significant correlation between intensity and time, as determined by the Spearman’s rank correlation test. Spearman’s correlation coefficients are shown next to the corresponding orange and blue dots at the top right of each panel.

Approaching response onset, neural information flow increased bidirectionally between the left and right precentral gyri (Spearman’s correlation coefficients: +0.73 and +0.70; p-values: 1.9 × 10^-5^ and 6.0 × 10^-5^) and between the left and right superior-frontal gyri (SFG) (Spearman’s correlation coefficients: +0.69 and +0.63; p-values: 1.1 × 10^-4^ and 6.3 × 10^-4^). After response onset, neural information flow decreased bidirectionally between these same regions (Spearman’s correlation coefficients: −0.84 to −0.57; p-values: 2.3 × 10^-5^ to 0.002).

### 3.6. Relationship between high-gamma modulation and behavioral performance

Logistic and linear mixed model analyses identified the timing and ROIs where high-gamma amplitudes were associated with response accuracy and response time, independent of the four fixed-effect covariates mentioned above. These analyses provided evidence that stronger neural engagement in the right aMFG enhanced response accuracy but delayed the response. Specifically, increase in high-gamma amplitude in the right aMFG between 400 and 500 ms post-stimulus onset was associated with higher odds of a successful response (odds ratio: 1.006; 95% CI: 1.003 to 1.010; maximum t-value: +3.6; p-value: 3.6 × 10^-4^; DF: 4109.0; Figure 6A). Increase in high-gamma amplitude in the right aMFG between 250 and 500 ms post-stimulus onset was associated with a delayed response (mixed model estimate: +0.9 ms/%; 95% CI: +0.4 to +1.5; maximum t-value: +3.4; p-value: 5.7 × 10^-4^; DF: 4037.0; Figure 6B), whereas stronger high-gamma augmentation in the left precentral gyrus between 250 and 500 ms post-stimulus onset was associated with a faster response (mixed model estimate: −1.0 ms/%; 95% CI: −1.4 to −0.5; minimum t-value: −3.9; p-value: 9.4 × 10^-5^; DF: 7073.0).

**Figure 6.**
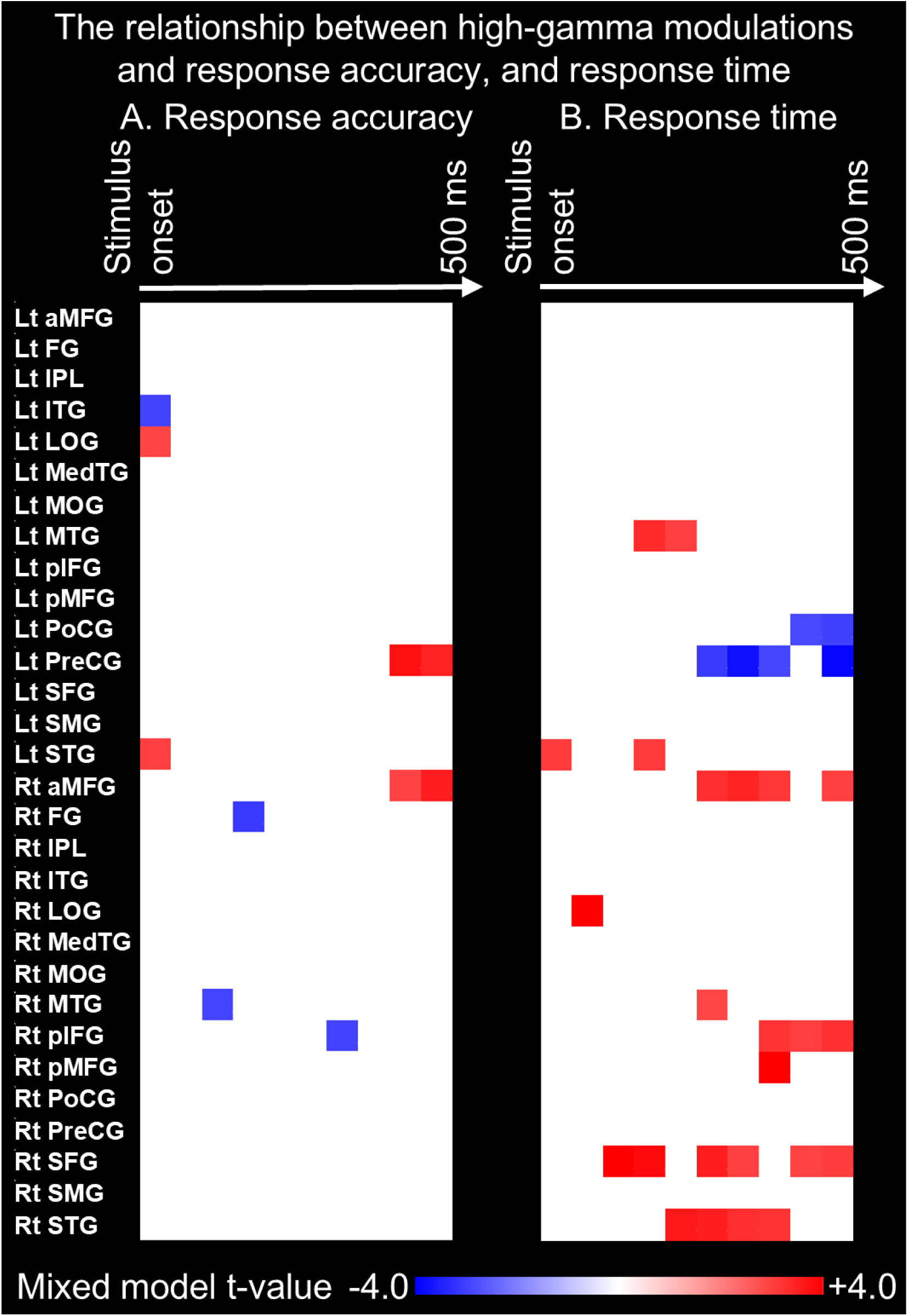
Relationship between high-gamma modulations and response accuracy, and response time. The matrices display mixed model t-values, representing the strength of association with (A) response accuracy and (B) response time in each region of interest during every 50-ms time window. Refer to Figure 1 for the definitions of each abbreviation.

The high-gamma effect, which prolonged response time with significance, was also noted in extensive regions within the right hemisphere, including SFG, posterior middle-frontal gyrus (pMFG), posterior inferior-frontal gyrus, superior-temporal gyrus, middle-temporal gyrus between 250 and 500 ms post-stimulus period (mixed model estimates: +0.6 to +1.1 ms/%; maximum t-values: +2.9 to +4.7; p-values: 3.0 × 10^-6^ to 0.003; Figure 6B). The right aMFG is directly connected with these frontal ROIs via local u-fibers.

## 4. Discussion

### 4.1. Significance and innovation

This study is the first to present a dynamic tractography map visualizing millisecond-scale functional connectivity and neural information flow between distant cortical lobes via specific white matter tracts during visuomotor integration. Intracranial recordings of high-gamma activity enabled per-trial assessments of task-related neural engagement at millisecond temporal resolution and allowed evaluation of its impact on behavioral performance of given participants. By integrating group-level neural dynamics recorded from distant cortical regions directly connected by white matter pathways (Kanno et al., 2025), we inferred biologically plausible neural communication via specific fasciculi.

Our observations clarify the neural communication dynamics underlying visuomotor integration. Specifically, visuospatial information is initially perceived in occipital regions and subsequently transferred to temporal regions via the inferior longitudinal fasciculus. Further visuomotor transformations—converting visuospatial representations into motor commands— occur predominantly within the right hemisphere across association cortices through white matter bundles, including the superior longitudinal and arcuate fasciculi. Concurrently, neural activity in the right aMFG supports cognitive control, facilitating appropriate response selection during periods requiring additional deliberation and continuous response monitoring. Finally, the corpus callosum conveys these decisions—primarily generated within right-hemispheric networks—to the left-hemispheric motor cortex approximately 100 ms before action execution. Collectively, these findings address a critical knowledge gap regarding how the right hemisphere coordinates visual inputs and how the left hemisphere subsequently executes corresponding motor actions.

### 4.2. Neural communication for visuospatial perception and analysis

Neural communication underlying visuospatial perception and analysis is demonstrated by local high-gamma modulations following stimulus onset (**Video S2**), as well as functional connectivity enhancements and neural information flows (**Videos S3 and S4**). Occipital high-gamma augmentation became evident by 120 ms after stimulus onset. Enhanced functional connectivity through the inferior longitudinal fasciculus, linking occipital regions to the fusiform and medial temporal regions, emerged 200–300 ms after stimulus onset, accompanied by occipital-to-temporal neural information flows. Concurrently, inter-hemispheric connectivity enhancement via callosal fibers between bilateral occipital lobes was observed, characterized by left-to-right neural information flows.

Local high-gamma augmentation in occipital regions may represent perceptual processing of lower-order visual information and increased attentional modulation (**Video S2**). Supporting this interpretation, an iEEG study demonstrated that full-field flash visual stimuli elicited high-gamma augmentation in medial and lateral occipital regions approximately 100 ms post-stimulus, and electrical stimulation at these sites frequently induced phosphene sensations (Nakai et al., 2018). Another intracranial study reported that sustained attention to visual stimuli further enhanced high-gamma augmentation in the lateral occipital region (Tallon-Baudry et al., 2005).

High-gamma augmentation in fusiform and medial temporal regions, along with enhanced functional connectivity between occipital and temporal areas (**Video S3**), likely reflects perceptual processing of higher-order visual information. While passive viewing of full-field flash stimuli did not elicit high-gamma augmentation in the fusiform regions (Nakai et al., 2018), attention to simple or common objects or written letters has been reported to elicit robust high-gamma augmentation in fusiform regions (Tallon-Baudry et al., 2005; Thesen et al., 2012; Nakai et al., 2019). Moreover, high-gamma responses in the fusiform region exhibit strong selectivity among individual exemplars within the category of faces, an effect that cannot be explained by differences in low-level visual features (Davidesco et al., 2014). Electrical stimulation of the fusiform gyrus frequently elicits visual distortions, such as the perception of objects or altered visual images (Parvizi et al., 2012; Rangarajan et al., 2014; Nakai et al., 2018). fMRI studies have demonstrated that face stimuli preferentially elicit hemodynamic activation within the fusiform gyrus, whereas building or scene stimuli selectively activate the parahippocampal gyrus, a region situated in the medial temporal cortex (Epstein et al., 1999; Kanwisher and Yovel, 2006). Another fMRI study reported enhanced activation in the parahippocampal gyrus specifically in response to visual stimuli characterized by sharp edges (Rajimehr et al., 2011), similar to the visual stimuli used in the present study (Figure 2). Additionally, electrical stimulation of the parahippocampal gyrus has been shown to induce topographic visual hallucinations (e.g., seeing indoor or outdoor scenes) and transient memory impairments (Mégevand et al., 2014; Tani et al., 2016).

Enhanced inter-hemispheric functional connectivity and left-to-right neural information flow between occipital lobes (**Video S4**) likely reflect the integration of bilateral visual inputs, facilitating the combination of visual information from the right and left visual fields into a unified perceptual experience. Increased left-to-right information transfer may represent mechanisms by which the right hemisphere integrates low-order visual percepts initially processed in the left hemisphere, thereby supporting comprehensive spatial perception. Previous studies have reported that corpus callosotomy can impair recognition of objects explicitly presented to the left visual field, a deficit attributed to disrupted inter-hemispheric transfer of perceptual representations (Lausberg et al., 2003; Bloom and Hynd, 2005; Gazzaniga, 2005). Moreover, impaired object recognition following callosotomy is particularly pronounced when the posterior portion of the corpus callosum is disconnected (Kawai et al., 2004).

### 4.3. Neural communication for visuomotor transformation and cognitive control

The present study identified enhanced functional connectivity and neural information flows, likely reflecting processes involved in visuomotor transformation. Specifically, between 250 ms post-stimulus and 100 ms before response execution, neural information flowed bidirectionally through the right superior longitudinal fasciculus, connecting frontal regions (e.g., MFG and precentral gyrus) with parietal regions (supramarginal gyrus and inferior parietal lobule; **Video S4**). Concurrently, neural information was transferred through the right arcuate fasciculus from the middle-temporal gyrus to the aMFG and precentral gyrus. Lesion studies have shown that damage to the right superior longitudinal and arcuate fasciculi frequently impairs imitation of hand postures, attributed either to disruptions in visuomotor transformation or visuospatial neglect (Leiguarda and Marsden, 2000; Dressing et al., 2020). Furthermore, repetitive transcranial magnetic stimulation (rTMS) studies in healthy adults have demonstrated that virtual lesions of the right parietal lobe can impair visual stimulus detection (Hilgetag et al., 2001; Rounis et al., 2007). Another rTMS study indicated that stimulation of the right parietal cortex, both within the initial 500 ms post-stimulus and during an additional subsequent 500 ms interval, additively prolonged reaction times in a visual search task (Rosenthal et al., 2006).

The present study also identified enhanced functional connectivity and local bidirectional information exchange among the right aMFG, pMFG, and precentral gyrus via short-range u-fibers, likely reflecting cognitive control processes. Specifically, at 400–500 ms post-stimulus onset, each 1% increase in high-gamma augmentation in the right aMFG was associated with a 0.6% increase in the odds of producing a successful response, accompanied by a 0.9 ms increase in response latency (Figure 6). fMRI studies of healthy adults have shown that a nonverbal visuomotor task requiring conflict monitoring elicits enhanced hemodynamic activation in the right dorsolateral prefrontal region, including the aMFG (van Gaal et al., 2010; Yang et al., 2024). Studies of patients with brain tumors reported that intraoperative electrical stimulation of cortical and white matter regions at the level of right MFG and inferior frontal gyrus frequently resulted in selective impairment in cognitive control, as evidenced by failed performance on a Stroop task (Puglisi et al., 2018; 2019). Consistent with these findings, our previous iEEG study demonstrated that high-gamma amplitude in the right aMFG was specifically enhanced approximately 400 ms prior to responses during a Stroop task (Sakakura et al., 2024). Other iEEG studies using go/no-go tasks similarly reported increased high-gamma amplitude in the right aMFG following stop signals (Swann et al., 2013; Fonken et al., 2016). Additionally, iEEG investigations employing visuomotor tasks observed enhanced high-gamma amplitude in the right aMFG immediately after erroneous responses (Völker et al., 2018; Mitsuhashi et al., 2022).

### 4.4. Neural communication for motor responses

The present study clarified the dynamics of neural communication underlying motor execution in the left precentral gyrus as driven by decisions made in the right hemisphere. Specifically, approximately 100 ms before response onset, neural information flowed bidirectionally through the corpus callosum between the right and left precentral gyri (**Video S4**). Around the time of response onset, high-gamma augmentation peaked in the left precentral gyrus (**Video S2**). Prior intracranial studies have demonstrated that frontal cortices can deliver neural information directly to their homotopic counterparts in the contralateral hemisphere through the corpus callosum within 30 ms, based on the latency of cortical responses to single-pulse electrical stimulation (Trebaul et al., 2018; Mitsuhashi et al., 2021). Moreover, studies in healthy adults have shown that TMS of the motor cortex can induce either facilitation or inhibition of the contralateral motor cortex (Ferbert et al., 1992; Hanajima et al., 2001; Daskalakis et al., 2002; Bortoletto et al., 2021). Additionally, callosotomy has been reported to result in akinetic mutism, characterized by an inability to initiate movement (Crutchfield et al., 1994). iEEG studies further suggest that cortical sites exhibiting high-gamma augmentation during motor responses are far more likely to produce motor symptoms when electrically stimulated (Nakai et al., 2017; Arya et al., 2018).

### 4.5. Methodological considerations

The spatial extent of intracranial electrode placement is determined solely by clinical necessity; therefore, the spatial coverage of iEEG signal sampling from non-epileptic areas is inherently limited for each patient. To address this spatial sampling constraint, previous iEEG studies have aggregated data across multiple patients, using group-averaged high-gamma amplitudes to estimate neural communication dynamics representative of the broader population (Kunii et al., 2013; Burke et al., 2014; Avanzini et al., 2016; Solomon et al., 2017; Kanno et al., 2025). In other words, our study was not designed to localize neural communication pathways at the individual patient level.

Previously, we validated the methodological approach of using group-averaged high-gamma amplitudes to quantify functional connectivity enhancements during a language task (Kanno et al., 2025). Specifically, we demonstrated that cortical areas exhibiting greater functional connectivity enhancements were more likely to elicit receptive and expressive aphasia upon electrical stimulation. Importantly, the association between functional connectivity enhancements and stimulation-induced language deficits was stronger than that between local high-gamma augmentation alone and stimulation-induced language deficits (Kanno et al., 2025).

We acknowledge that within ROIs containing more electrode sites, the influence of individual electrodes on group averages is comparatively reduced. For example, the left and right precentral gyri contained 41 and 46 electrode sites, respectively, whereas the left and right aMFG had only 14 and 26 sites (**Table S1**). Consequently, functional connectivity measures derived from precentral ROIs inherently benefit from approximately 50% higher signal-to-noise ratios compared to measures from aMFG ROIs. However, since our analysis incorporated trial-level high-gamma measures—with each patient completing an average of 184 trials—we maintained sufficient statistical power to detect even a small effect size (Cohen’s d ≈ 0.20) in high-gamma amplitude modulation, even within ROIs containing as few as three electrode sites.

Subsets of white matter pathways exhibiting significant enhancement in functional connectivity did not demonstrate facilitatory neural information flows, an observation warranting cautious interpretation. The algorithm described in Ito et al. (2011), which computes neural information flow based on relative timing of high-gamma amplitude modulations between two ROIs, inherently requires sufficient data points for reliable analysis. Increasing the number of time points improves the power to detect consistent temporal patterns of high-gamma amplitude changes at one ROI preceding those at another ROI. However, the assumption of stationary neural information flow becomes progressively less tenable for analysis periods substantially longer than 250 ms. Given these technical constraints, our analysis primarily aimed to clarify the directionality of neural information flows during periods of enhanced functional connectivity in a visuomotor task. We found neural information flow from the occipital to temporal lobe and from the left to right occipital lobe following stimulus onset, whereas neural information flow between the right and left precentral gyri around response onset was bidirectional (**Video S3**).

Behavioral studies suggest that the speed of visuomotor integration typically increases markedly from school age (approximately 6–10 years) through adolescence and young adulthood, before gradually declining in older adulthood (around 75 years) (Liu et al., 2014; Richards et al., 2017). In the present study, involving patients aged 9 to 20 years, we did not observe a significant effect of age on behavioral performance or high-gamma measures. This finding should be interpreted cautiously, as our relatively narrow age range and modest sample size may have limited statistical power to detect subtle developmental effects on the neural communication pathways supporting visuomotor integration. Rather, our study specifically aimed to delineate the dynamics of neural communication through major white matter pathways consistently observed across the general population.

## Acknowledgements

We are grateful to Sandeep Sood, MD, Neena I. Marupudi, MD, and Jamie MacDougall, RN, BSN, CPN at Children’s Hospital of Michigan, for the collaboration and assistance in performing the studies described above. We want to thank Lumos Labs for providing us with the software as a part of the Lumosity Human Cognition Project (https://www.lumosity.com/hcp).

## CRediT authorship contribution statement

**Riyo Ueda:** Conceptualization, Data curation, Formal analysis, Investigation, Methodology, Software, Validation, Visualization, Writing – original draft. **Hiroshi Uda:** Data curation, Validation, Visualization, Writing – review and editing. **Keisuke Hatano:** Validation, Visualization, Writing – review and editing. **Kazuki Sakakura:** Data curation, Formal analysis, Methodology, Software. **Naoto Kuroda:** Writing – review and editing. **Yu Kitazawa:** Software. **Aya Kanno:** Data curation. **Min-Hee Lee:** Software. **Jeong-Wong Jeong:** Software. Resources. **Aimee F. Luat:** Data curation. **Eishi Asano:** Conceptualization, Data curation, Funding acquisition, Investigation, Methodology, Resources, Supervision, Validation, Writing – review and editing.

