## Supplementary document for "Millisecond-Scale White Matter Dynamics Underlying Visuomotor Integration"

This document includes:
**Tables S1-S4**;
Legends for **Videos S1-S4.**

| Regions of interest | Electrodes | Patients |
|---|---|---|
| Lt aMFG: left anterior middle-frontal gyrus | 14 | 3 |
| Lt FG: left fusiform gyrus | 18 | 4 |
| Lt IPL: left inferior parietal lobule | 9 | 3 |
| Lt ITG: left inferior-temporal gyrus | 16 | 4 |
| Lt LOG: left lateral occipital gyrus | 10 | 3 |
| Lt MedTG: left medial temporal gyrus (entorhinal and parahippocampal gyri) | 4 | 3 |
| Lt MOG: left medial occipital gyrus (lingual and cuneus gyri) | 3 | 2 |
| Lt MTG: left middle-temporal gyrus | 23 | 3 |
| Lt pIFG: left posterior inferior-frontal gyrus (BA 44 and 45) | 7 | 2 |
| Lt pMFG: left posterior middle-frontal gyrus | 14 | 4 |
| Lt PoCG: left postcentral gyrus | 32 | 4 |
| Lt PreCG: left precentral gyrus | 41 | 3 |
| Lt SFG: left superior frontal gyrus | 19 | 1 |
| Lt SMG: left supramarginal gyrus | 30 | 3 |
| Lt STG: left superior-temporal gyrus | 34 | 4 |
| Rt aMFG: anterior middle-frontal gyrus | 26 | 4 |
| Rt FG: fusiform gyrus | 26 | 4 |
| Rt IPL: inferior parietal lobule | 19 | 4 |
| Rt ITG: inferior-temporal gyrus | 19 | 4 |
| Rt LOG: lateral occipital gyrus | 19 | 4 |
| Rt MedTG: medial temporal gyrus (entorhinal and parahippocampal gyri) | 19 | 4 |
| Rt MOG: medial occipital gyrus (lingual and cuneus gyri) | 17 | 4 |
| Rt MTG: middle-temporal gyrus | 20 | 3 |
| Rt pIFG: posterior inferior-frontal gyrus (BA 44 and 45) | 19 | 4 |
| Rt pMFG: posterior middle-frontal gyrus | 21 | 5 |
| Rt PoCG: postcentral gyrus | 24 | 4 |
| Rt PreCG: precentral gyrus | 46 | 4 |
| Rt SFG: superior frontal gyrus | 30 | 5 |
| Rt SMG: supramarginal gyrus | 29 | 4 |
| Rt STG: superior-temporal gyrus | 29 | 4 |
| **Total** | 637 | 8 |

**Table S1. The number of artifact-free, nonepileptic electrode sites used for region of interest (ROI)-based analysis.** All aforementioned ROIs (15 per hemisphere) were included in the ROI-based analysis, with each containing at least three electrodes. Lt: left. Rt: right.

| **Regions of interest** | **Electrodes** | **Patients** |
|---|---|---|
| Lt aCG: anterior cingulate gyrus (rostral and caudal anterior cingulate) | 0 | 0 |
| Lt FP: frontal pole | 0 | 0 |
| Lt OrbF: left orbitofrontal region (BA 11, 12 and 47) | 0 | 0 |
| Lt pCG: posterior cingulate gyrus (posterior and isthmus cingulate) | 0 | 0 |
| Lt PCL: paracentral lobule | 2 | 1 |
| Lt PCun: precuneus | 0 | 0 |
| Lt SPL: superior parietal lobule | 0 | 0 |
| Rt aCG: anterior cingulate gyrus (rostral and caudal anterior cingulate) | 0 | 0 |
| Rt FP: frontal pole | 2 | 1 |
| Rt OrbF: orbitofrontal region (BA 11, 12 and 47) | 0 | 0 |
| Rt pCG: posterior cingulate gyrus (posterior and isthmus cingulate) | 7 | 3 |
| Rt PCL: paracentral lobule | 3 | 1 |
| Rt PCun: precuneus | 10 | 2 |
| Rt SPL: superior parietal lobule | 9 | 2 |
| **Total** | 33 | 4 |

**Table S2. Regions excluded from region-of-interest (ROI)-based analysis.** Regions were excluded from ROI-based analysis because fewer than three electrode sites were available within an ROI in the left hemisphere. Intracranial EEG signals from these ROIs were included in the map illustrating task-related high-gamma amplitude modulations (**Video S2**).

| Parameters | Coefficient [ms] | DF | t-value | p-value | 95% CI | |
|---|---|---|---|---|---|---|
| | | | | | Lower bound | Upper bound |
| Intercept | +839 | 5.2 | +2.6 | 0.048 | +11 | +1667 |
| Different symbol | +276 | 1227.2 | +9.0 | $7.3 \times 10^{-19}$ | +216 | +336 |
| Task switch | -33 | 1227.2 | -1.1 | 0.28 | -93 | +27 |
| Task familiarity | -2 | 1231.6 | -7.4 | $2.0 \times 10^{-13}$ | -3 | -1 |
| Prior failure | +306 | 1228.4 | +5.9 | $3.6 \times 10^{-9}$ | +205 | +407 |
| Age | +35 | 5.0 | +1.7 | 0.16 | -20 | +90 |

**Table S3. The relationship between predictor variables and response time.** CI, confidence interval; DF, degree of freedom.

| Parameters | DF | Coefficient | t-value | p-value | Odds ratio | 95% CI | |
|---|---|---|---|---|---|---|---|
| | | | | | | Lower bound | Upper bound |
| Intercept | 1257.0 | 1.549 | 1.2 | 0.24 | 4.709 | 0.364 | 60.875 |
| Different symbol | 1257.0 | -0.435 | -2.2 | 0.026 | 0.647 | 0.442 | 0.948 |
| Task switch | 1257.0 | -0.016 | -0.1 | 0.93 | 0.984 | 0.675 | 1.434 |
| Task familiarity | 1257.0 | $-6.3 \times 10^{-4}$ | -0.4 | 0.71 | 0.999 | 0.996 | 1.003 |
| Prior failure | 1257.0 | -0.912 | -3.7 | $2.1 \times 10^{-4}$ | 0.402 | 0.248 | 0.651 |
| Age | 1257.0 | 0.079 | +1.7 | 0.092 | 1.082 | 0.987 | 1.185 |

**Table S4. The relationship between predictor variables and response accuracy.** CI, confidence interval; DF, degree of freedom.

**Video S1. Visuomotor task.** The first author (R.U.) demonstrates the *Speed Match* game on the Lumosity platform, illustrating the visuomotor task used in this study.

**Video S2. Task-related high-gamma amplitude modulations.** This video illustrates the percentage changes in task-related high-gamma amplitudes relative to the mean amplitude during the entire 750-ms analysis period.

**Video S3. Functional connectivity and neural information flow.** This video illustrates the evolution of functional connectivity and neural information flow across three 250-ms time windows around stimulus and response onsets.

**Video S4. Spatiotemporal dynamics of functional connectivity and neural information flow.** Color-coded streamlines indicate different types of functional connectivity modulations, while moving spheres depict the direction and intensity of neural information flow.
